# The human-specific paralogs SRGAP2B and SRGAP2C differentially modulate SRGAP2A-dependent synaptic development

**DOI:** 10.1101/596940

**Authors:** Ewoud R.E. Schmidt, Justine V. Kupferman, Michelle Stackmann, Franck Polleux

**Author notes:** These authors contributed equally (Ewoud R.E. Schmidt, Justine V. Kupferman). Address correspondence to: Franck Polleux, Ph.D., Columbia University, Department of Neuroscience, Mortimer B. Zuckerman Mind Brain Behavior Institute, Kavli Institute for Brain Science, Jerome L. Greene Science Center, 3227 Broadway, L-5-050, MC 9853, New York, NY 10027.

## Abstract

Human-specific gene duplications (HSGD) have recently emerged as key modifiers of brain development and evolution. However, the molecular mechanisms underlying the function of HSGDs remain often poorly understood. In humans, a truncated duplication of SRGAP2A led to the emergence of two human-specific paralogs: SRGAP2B and SRGAP2C. The ancestral copy SRGAP2A limits synaptic density and promotes maturation of both excitatory (E) and inhibitory (I) synapses received by cortical pyramidal neurons (PNs). SRGAP2C binds to and inhibits all known functions of SRGAP2A leading to an increase in E and I synapse density and protracted synapse maturation, traits characterizing human cortical neurons. Here, we demonstrate how the evolutionary changes that led to the emergence of SRGAP2 HSGD generated proteins that, in neurons, are intrinsically unstable and upon hetero-dimerization with SRGAP2A, reduces SRGAP2A levels in a proteasome-dependent manner. Moreover, we show that, compared to SRGAP2B, and despite only a few non-synonymous mutations specifically targeting arginine residues, SRGAP2C is unique, compared to SRGAP2B, in its ability to induce long-lasting changes in synaptic density throughout adulthood. The non-synonymous mutations specifically targeting arginine residues led to the unique ability of SRGAP2C to inhibit SRGAP2A function and thereby contribute to the emergence of human-specific features of synaptic development during evolution.

## Introduction

Evolutionary changes that occurred specifically in the human genome have been proposed to be significant drivers of human speciation. Recently, significant progress has been made in cataloging human-specific genetic changes, especially with respect to evolution of the human brain [1]. However, knowledge of how these human-specific genetic modifications impact corresponding molecular pathways is still lacking. For example, multiple human-specific gene duplications (HSGD) have recently been identified to play a role in controlling human brain development [2–11]. However, how these genetic modifiers affect specific signaling pathways in the developing or adult human brain remains poorly understood. This limits our understanding of how these changes led to the emergence of human-specific biological traits.

Previous work has demonstrated how gene duplication of SLIT-ROBO Rho-GTPase-activating protein 2 (SRGAP2A), which is highly expressed in the developing brain of all mammals [12,13], resulted in emergence of the human-specific paralogs SRGAP2B and SRGAP2C [8,14]. During human evolution, large segmental duplication of SRGAP2A led to the sequential emergence of SRGAP2B and SRGAP2C. Both are truncated open reading frames (only first 9 exons out of 22 in *Srgap2a*) leading to the expression of a truncated protein corresponding to the F-BAR domain, missing the last C-terminal 49 amino acids (F-BAR^Δ49^) (Fig. 1B). In addition, SRGAP2C acquired a series of unique nonsynonymous base pair mutations selectively targeting five arginine residues compared to SRGAP2B [6,8] (Fig. 1C). This truncation and these specific arginine mutations have been shown to reduce solubility of SRGAP2C and increase its ability to heterodimerize with SRGAP2A to form an insoluble complex [15].

**Fig. 1.**
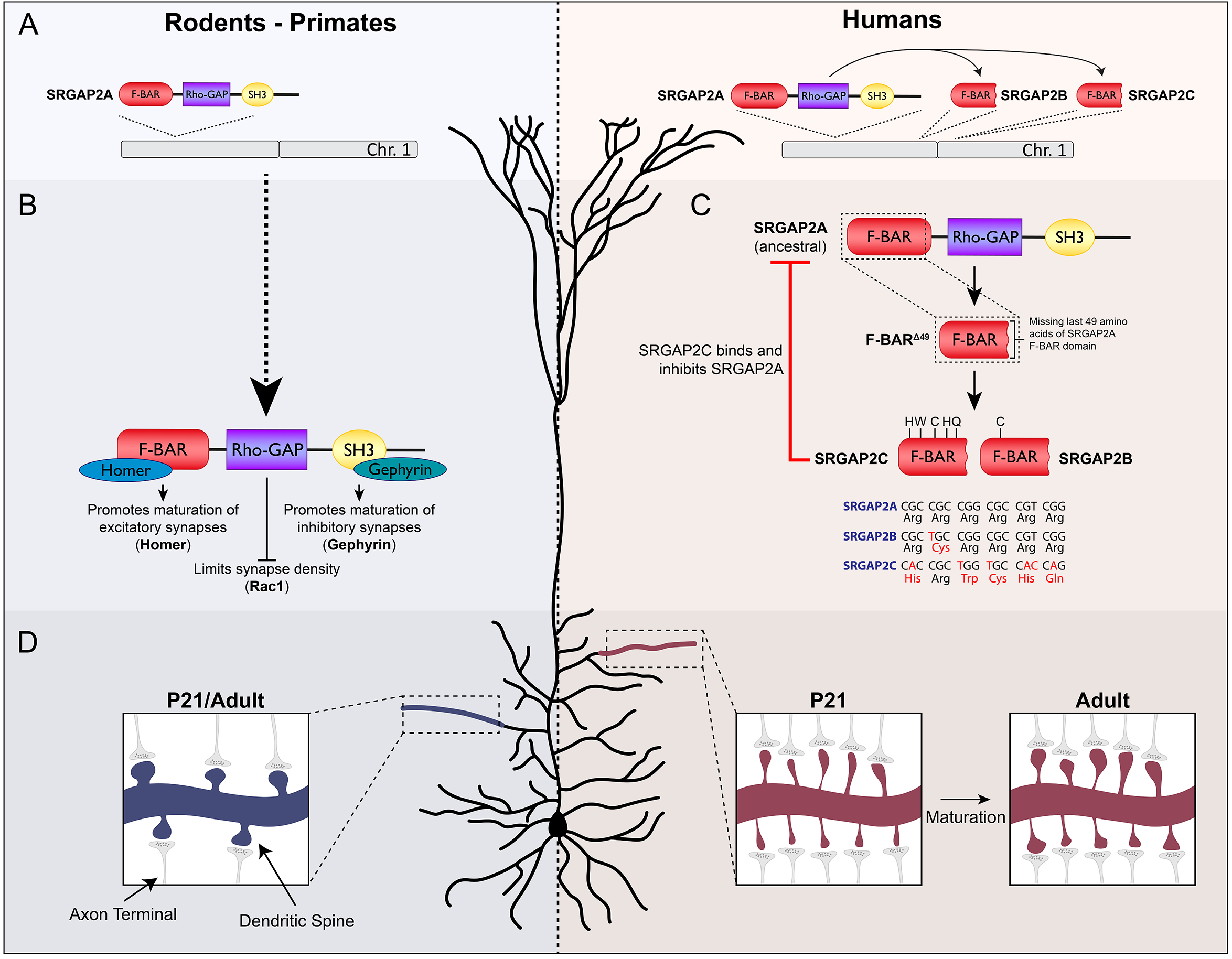
SRGAP2A and the emergence of the human-specific paralog SRGAP2C. (**A**) The ancestral copy SRGAP2A, present in rodents, primates, and humans, was duplicated in the human lineage to form the truncated copies SRGAP2B and SRGAP2C. (**B**) SRGAP2A contains three distinct protein domains. Through binding of Homer to an EVH1 site in the F-BAR domain, SRGAP2A promotes maturation of excitatory synapses, while binding of Gephyrin to the SH3 domain promotes inhibitory synapse maturation. The Rho-GAP domain is involved in limiting synapse density through Rac1. (**C**) Duplication of SRGAP2A in humans generated a truncated protein, F-BAR^Δ49^, containing the F-BAR domain of SRGAP2A that lacks the last C-terminal 49 amino acids. Subsequent nonsynonymous base pair mutations converted arginines to non-polar residues which led to the formation of SRGAP2B and SRGAP2C. Binding of SRGAP2C to SRGAP2A inhibits all functions of SRGAP2A. (**D**) In rodents, synaptic development of cortical pyramidal neurons is complete at P21. At this stage, both synapse size and synaptic density reaches levels of those observed in adulthood. In contrast, expression of SRGAP2C in mouse cortical pyramidal neurons binds and inhibits SRGAP2A resulting in protracted synaptic maturation and increased synaptic density.

Interestingly, expression of SRGAP2C in mouse cortical pyramidal neurons leads to the emergence of human-specific traits of synaptic development, including an increase in synaptic density and protracted synaptic maturation of both excitatory and inhibitory synapses [6,7]. This phenotype is strikingly similar to that of a constitutive knockout mouse in which *Srgap2a* expression is genomically reduced, strongly suggesting that human-specific SRGAP2C functions largely by inhibiting SRGAP2A function (Charrier et al. 2012). Interestingly, the ancestral copy SRGAP2A limits E and I synapse density through its Rac1-specific GAP domain, while promoting maturation of both E and I synapses through its ability to bind to the postsynaptic scaffolding protein Homer1 at E synapses through its class II EVH1 binding domain embedded in its F-BAR domain, and the postsynaptic protein Gephyrin through its SH3 domain at I synapses, respectively [6,7,16,17] (Fig 1A). Because SRGAP2C directly binds SRGAP2A through its truncated F-BAR domain, we previously hypothesized that this binding directly inhibits the function of SRGAP2A [6]. However, the mechanisms underlying the ability of SRGAP2C to inhibit all functions of SRGAP2A remained unknown.

Recent evidence suggests that both SRGAP2B and SRGAP2C are expressed in the human brain [2]. In the human population, *SRGAP2C* copy numbers are remarkably fixed, while *SRGAP2B* exhibits significant copy number variation (CNV) in the human population [8]. This suggests that in contrast to SRGAP2B, SRGAP2C has been rapidly fixed in the human population since its emergence during human brain evolution, and suggests that SRGAP2C might have played a unique role compared to SRGAP2B. However, the human-specific paralog SRGAP2B has not been characterized functionally but is almost identical to SRGAP2C in all other respects. SRGAP2B also lacks the last 49 amino acids of its F-BAR domain, but has its own unique combination of point mutations that led to five arginine mutations into non-polar residues in SRGAP2C (Dennis et al. 2012; Charrier et al. 2012).

Here we provide evidence showing that in cortical neurons, the truncation of the F-BAR domain present in both SRGAP2B and SRGAP2C leads to proteasome-mediated degradation of these proteins. Moreover, upon binding to SRGAP2A, SRGAP2C targets this hetero-dimer to the proteasome degradation pathway, thereby effectively reducing SRGAP2A protein levels in dendrites of cortical PNs. We show that SRGAP2C is uniquely more potent than SRGAP2B at long-lasting increase of synaptic density into adult cortical PNs. Together, these results show how the emergence of the human-specific paralog SRGAP2C directly impacted the ancestral copy SRGAP2A, a critical regulator of synaptic development during human brain evolution.

## Results

Human-specific partial duplication of *SRGAP2A* resulted in the emergence of two truncated paralogs (*SRGAP2B* and *SRGAP2C*) which both encode for a truncated protein consisting of the F-BAR domain of SRGAP2A lacking the last C-terminal 49 amino acids (Dennis et al. 2012; Charrier et al. 2012) (F-BAR^Δ49^; Fig 1). In addition, SRGAP2C acquired a series of point mutations that led to five non-synonymous amino acid changes that in SRGAP2A are all arginine residues (Fig 1). First, we examined the effects of the Δ49 truncation. Previous work using recombinant proteins and cell lines showed that this Δ49 truncation of the F-BAR domain generates intrinsically unstable and insoluble protein aggregates [15]. In order to investigate whether the Δ49 truncation caused the formation of similar aggregates in neurons, we co-expressed the full-length F-BAR domain of SRGAP2A (F-BAR), F-BAR with the Δ49 truncation (F-BAR^Δ49^), and SRGAP2C mRFP-tagged proteins together with Venus (cell fill) by *ex utero* electroporation in mouse cortical pyramidal neurons (PNs) cultured for 18-21 days *in vitro* (DIV) (Fig 2 and S1). Expression of F-BAR protein was clearly observed in soma and dendrites of cortical PNs. In contrast, F-BAR^Δ49^ and SRGAP2C expression levels were generally very low (Fig 2A and S2). We wondered whether the low expression of F-BAR^Δ49^ and SRGAP2C was the result of an active degradation mechanism, such as the proteasome degradation pathway, in response to the insolubility of these proteins. We therefore treated with MG-132, a potent and widely used inhibitor of proteasome function, and used a live-cell confocal imaging approach to measure protein levels in the same neurons over time. Upon treatment with MG-132, we observed a rapid increase of F-BAR^Δ49^ and SRGAP2C protein levels, while F-BAR levels increased only slightly (Fig 2A and B). These results show that the Δ49 truncation generates an insoluble protein that in neurons leads to proteasome-mediated degradation.

**Fig. 2.**
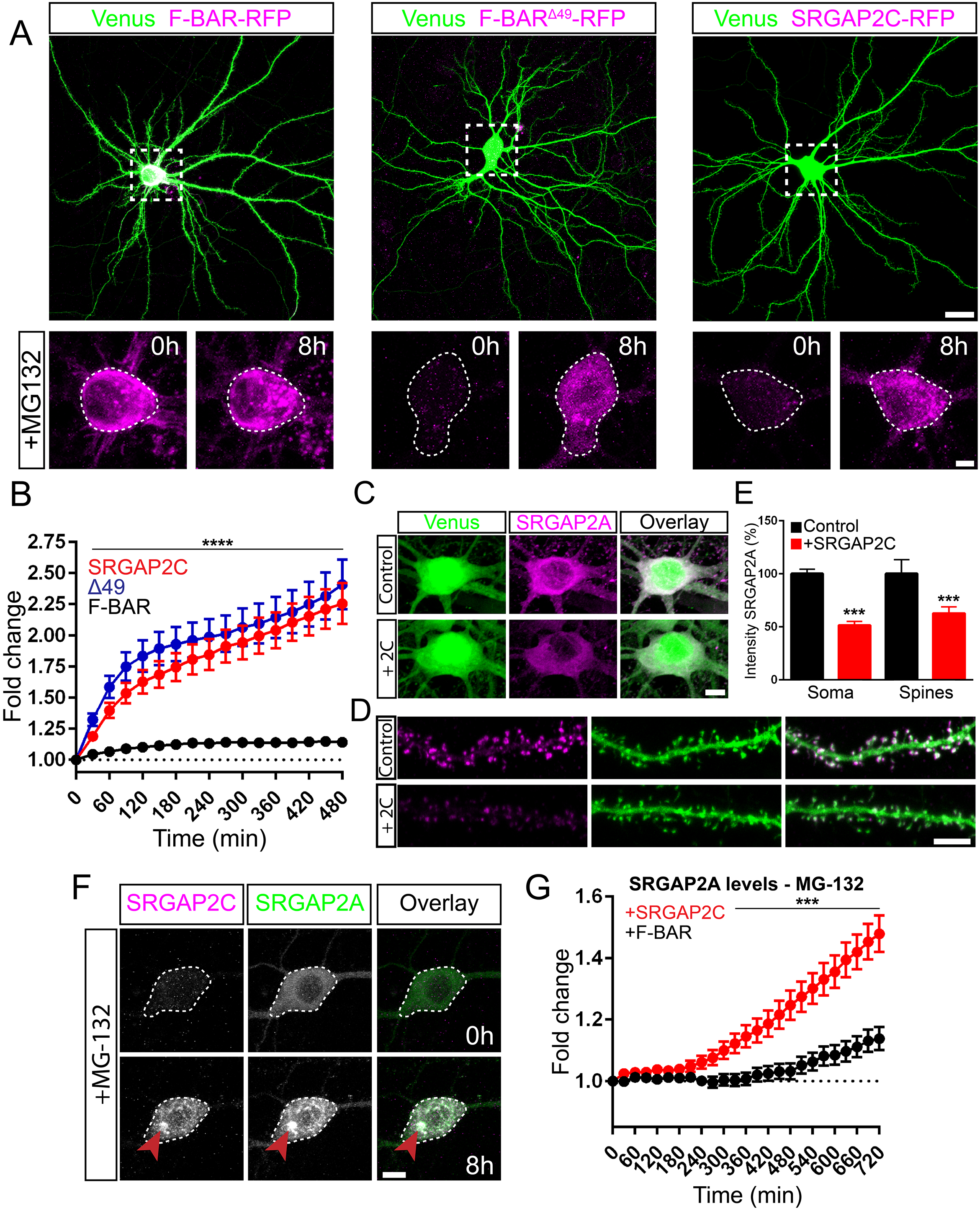
Proteasome degradation of SRGAP2 proteins. (**A, B**) Expression of RFP-tagged F-BAR, F-BAR^Δ49^, or SRGAP2C in mouse cortical pyramidal neurons cultured for 18-21 days *in vitro* (DIV). Low levels of expression were observed for F-BAR^Δ49^ and SRGAP2C, while addition of the proteasome inhibitor MG-132 resulted in a strong increase after 8h of treatment. Scale bar top panels: 25 µm, bottom panels: 5 µm. See (**B**) for quantification of fluorescence intensity after MG-132 treatment. n_F-BAR_ = 31, n_F-BAR_^Δ49^ = 40, n_SRGAP2C_ = 47; ****p < 0.0001; mean ± SEM. (**C–E**) Co-expression of SRGAP2A and SRGAP2C in cultured mouse cortical neurons. SRGAP2A levels are reduced when co-expressed with SRGAP2C in both soma and dendritic spines. Scale bars: 5 µm. See (**E**) for quantification of SRGAP2A fluorescence intensity. Soma: n_Control_ = 110, n_SRGAP2C_ = 68; Spines: n_Control_ = 969, n_SRGAP2C_ = 1259; ***p < 0.001; mean ± SEM. (**F**) Treatment of cultured mouse cortical neurons expressing both SRGAP2A-GFP and SRGAP2C-RFP with MG-132. Levels for both SRGAP2A and SRGAP2C increased upon treatment, and co-localization of SRGAP2A and SRGAP2C protein clusters were observed (red arrow). Scale bar: 10 µm. (**G**) Quantification of SRGAP2A fluorescent intensity when co-expressed with either F-BAR or SRGAP2C. n_F-BAR_ = 60, n_SRGAP2C_ = 59; ***p < 0.001; mean ± SEM.

Considering that SRGAP2C directly binds SRGAP2A, and that SRGAP2A plays a critical role in synaptic development [6,7], we next asked how SRGAP2C expression affects SRGAP2A protein. We co-expressed EGFP-tagged or mTagBFP-HA-tagged-SRGAP2A with or without mRFP-tagged SRGAP2C in mouse cortical neurons. Additionally, we co-expressed a cell fill, either Venus or mTagBFP-HA, for visualization of neuronal morphology. Interestingly, expression levels for SRGAP2A were significantly reduced in both soma (Fig 2C) and dendritic spines (Fig 2D) upon co-expression with SRGAP2C (Fig 2E).

Since SRGAP2C binds to SRGAP2A through their F-BAR domains (Charrier et al. 2012; Sporny et al., 2017), we next asked whether the reduction of SRGAP2A levels in response to ‘humanization’ of SRGAP2C expression is proteasome-dependent, i.e., if the mechanism leading to reduction of SRGAP2A function in presence of SRGAP2C is due to targeting of this heterodimer complex to the proteasome for degradation. We co-expressed SRGAP2A-GFP with either F-BAR-RFP or SRGAP2C-RFP and treated neurons with MG-132. While SRGAP2A levels were only slightly elevated upon treatment with MG-132 when co-expressed with the full-length F-BAR domain, we observed a significant increase when co-expressed with SRGAP2C (Fig 2F and G) and frequently observed the formation and co-localization of SRGAP2A and SRGAP2C protein clusters (Fig 2F).

We performed a series of similar experiments using the protein synthesis inhibitor Cycloheximide (Chx) to measure the rate of SRGAP2A protein turnover when co-expressed with SRGAP2C. Indeed, Chx treatment led to a reduction of protein fluorescence over time. Compared to F-BAR, co-expression with SRGAP2C significantly decreased the half-life of SRGAP2A (Fig. S3). Together, these observations show that the Δ49 truncation present in SRGAP2C leads to proteasome-mediated degradation of both SRGAP2C and SRGAP2A when co-expressed together.

Besides the Δ49 truncation, SRGAP2C also acquired a number of point mutations resulting in non-synonymous changes of five arginines. These mutations are highly specific to SRGAP2C (not present in SRGAP2B) and have most likely been under selective pressure over the course of human evolution [8]. Furthermore, both SRGAP2B and C are expressed during human brain development [2,8]. Finally, because SRGAP2B displays significantly more copy number variation (CNVs) in the human population than SRGAP2C, it has been proposed that these two HSGDs might have different functions and most importantly, might exert different strength in their ability to act as modifiers of SRGAP2A function. To test whether the arginine mutations specific to SRGAP2C contribute to its ability to regulate synaptic development differentially than SRGAP2B *in vivo*, we used *in utero* cortical electroporation (IUCE) to express Venus (cell filler) and mRFP-tagged fusion protein of either F-BAR, F-BAR^Δ49^, SRGAP2B, and SRGAP2C in layer 2/3 pyramidal cortical neurons *in vivo* (Fig. 3A and S4) and compare their ability to phenocopy a partial SRGAP2A loss-of-function (Charrier et al. 2012). We analyzed the ability of these proteins to regulate E synaptic density and maturation by quantifying spine density and spine head size of apical oblique dendrites at juvenile (P21 – Fig. 3) and adult (>P65 – Fig. 4) stages. Comparing the effect of SRGAP2C on synaptic development to F-BAR and F-BAR^Δ49^ allowed us to determine how the distinct evolutionary events, from the truncation of the F-BAR domain (present in both SRGAP2B and C) to the progressive acquisition of five arginine conversions (present only in SRGAP2C), contributed to human-specific evolution of synaptic development.

**Fig. 3.**
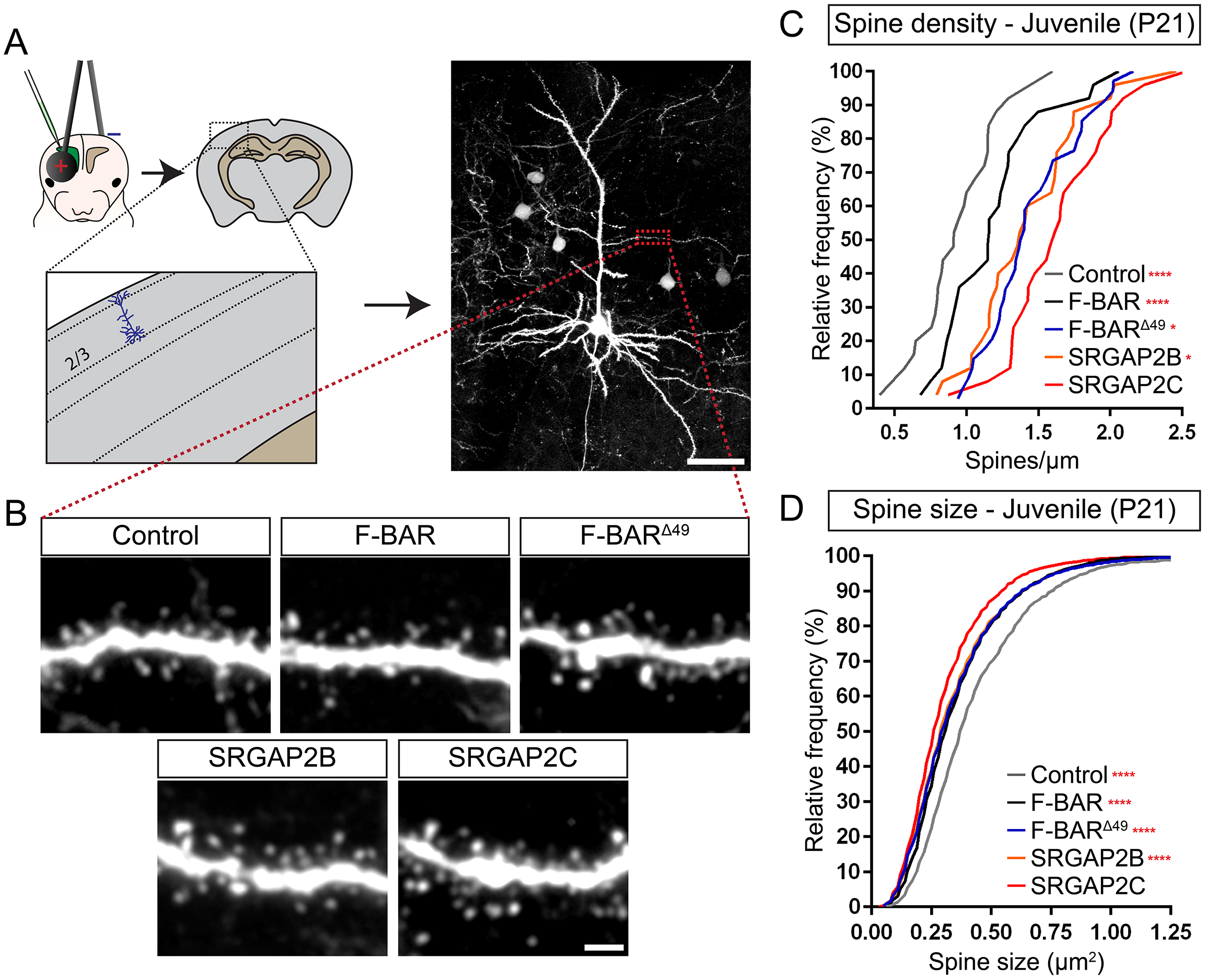
*In vivo* analysis of spine density and spine size at juvenile stage. (**A**) Schematic illustrating *in utero* electroporation approach targeting layer 2/3 cortical pyramidal neurons and subsequent quantification of dendritic spine density and head size. Scale bar: 50 µm. (**B**) Representative images showing the apical oblique dendrite of neurons at P21 co-expressing either Venus and tdTomato (Control), or mRFP-tagged F-BAR, F-BAR^Δ49^, SRGAP2B, or SRGAP2C. Scale bar: 2 µm. (**C**, **D**) Quantification of spine density (**C**) and spine head size (**D**). Quantifications are shown as frequency distributions. Spine density (segments): n_Control_ = 26, n_F-BAR_ = 25, n_F-BAR_^Δ49^ = 34, n_SRGAP2B_ = 25, n_SRGAP2C_ = 25; Spine size (number of spines): n_Control_ = 1693, n_F-BAR_ = 2054, n_F-BAR_^Δ49^ = 3267, n_SRGAP2B_ = 2900, n_SRGAP2C_ = 2448; *p < 0.05, ****p < 0.0001.

**Fig. 4.**
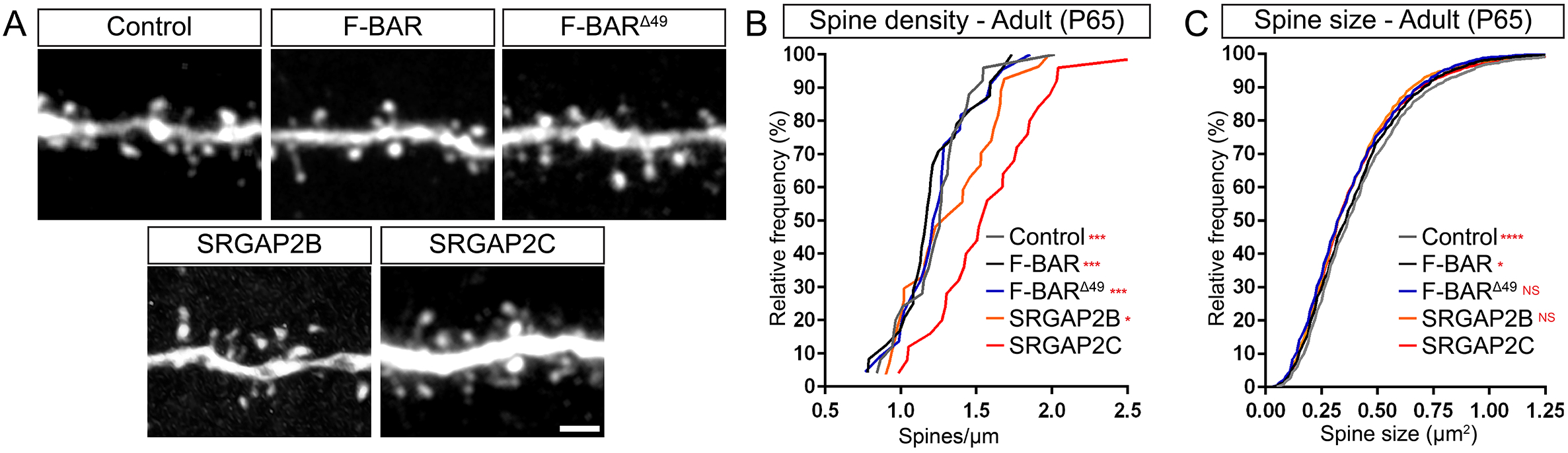
*In vivo* analysis of spine density and spine size at the adult stage. (**A**) Representative images showing the apical oblique dendrite of adult neurons co-expressing either Venus and tdTomato (Control), or mRFP-tagged F-BAR, F-BAR^Δ49^, SRGAP2B, or SRGAP2C. Scale bar: 2 µm. (**B, C**) Quantification of spine density (**B**) and spine head size (**C**). Quantifications are shown as frequency distributions. Spine density (segments): n_Control_ = 25, n_F-BAR_ = 24, n_F-BAR_^Δ49^ = 22, n_SRGAP2B_ = 27, n_SRGAP2C_ = 25; Spine size (number of spines): n_Control_ = 1872, n_F-BAR_ = 2049, n_F-BAR_^Δ49^ = 1843, n_SRGAP2B_ = 2295, n_SRGAP2C_ = 2630; NS: p > 0.05, *p < 0.05, ***p < 0.001, ****p < 0.0001.

We first analyzed how these proteins affected spine density. Compared to tdTomato controls, F-BAR slightly increased spine density at P21, while expression of SRGAP2C induced an increase in spine density at P21 that was significantly larger compared to either control or F-BAR expression (Fig. 3C). Interestingly, both F-BAR^Δ49^ and SRGAP2B had intermediary phenotypes, with increased spine density compared to control and F-BAR, but significantly smaller compared to SRGAP2C.

When analyzing spine head size, which is linearly proportional to postsynaptic AMPA receptor content and therefore classically used as an index of synaptic maturation [18–20], we found that for the control condition, spine head size at P21 was almost identical to that found at adult stages, showing that spines have reached maturity at P21 (Fig. 3D and 4C). In contrast, F-BAR expression led to reduced spine size compared to control at P21 (Fig. 3D). A similar reduction in spine size was observed for F-BAR^Δ49^ and SRGAP2B, while a significantly larger reduction in spine size was observed for SRGAP2C.

We next analyzed whether changes in spine density and head size observed at P21 persisted into adulthood for all the constructs tested (Fig. 4). Interestingly, only SRGAP2C significantly increased spine density in adults (Fig. 4A-B). The increased spine density observed for F-BAR^Δ49^ and SRGAP2B at P21 was no longer observed at adult stages. While spine head size was slightly smaller in adult animals for all of the proteins tested when compared to control, the difference in size was exceedingly small (for SRGAP2C a reduction of 8.2%), suggesting that for all conditions spines did mature to sizes which were very similar to controls at P65.

## Discussion

The first evolutionary step that led to the emergence of SRGAP2B and then SRGAP2C was an incomplete genomic duplication of the ancestral copy SRGAP2A generating the truncated protein F-BAR^Δ49^ [8,21]. This led to the formation of a highly insoluble protein that is sensitive to aggregate formation [15]. We show that in cortical pyramidal neurons this truncated protein is targeted to the proteasome for degradation. Moreover, our data also show that SRGAP2C reduces SRGAP2A levels in a proteasome-dependent manner. Since SRGAP2C directly binds SRGAP2A [6], our results demonstrate that SRGAP2C downregulates SRGAP2A function by binding to it and conferring the heterodimers proteosome-sensitivity. SRGAP2A plays a critical role in regulating synaptic density and maturation [6,7]. Restricting the available amount of SRGAP2A by targeting it to the proteasome represents an effective mechanism through which SRGAP2C increases synaptic density and delays maturation. Interestingly, HSGDs have often resulted in the formation of truncated paralogs and several of these truncated HSGDs have recently been implicated in regulating brain development [2–4,8,22,23]. This suggests that evolutionary changes affecting levels and patterns of expression not only occurred through modifications in regulators of gene expression at the genomic level [24], but also at the level of protein interactions through the emergence of insoluble proteins.

Following the initial duplication event of SRGAP2A, the newly formed SRGAP2B paralog acquired multiple base pair mutations specifically targeting five arginine residues, leading to the emergence of SRGAP2C. Previous work has shown how conversion of these arginine residues leads to reduced membrane binding for SRGAP2C and increased stability of the SRGAP2A:SRGAP2C heterodimer [15]. Besides SRGAP2C, the human-specific paralog SRGAP2B is also expressed in the human brain [2]. One of the question raised by this finding is whether SRGAP2B and SRGAP2C are redundant in their ability to downregulate SRGAP2A function or whether SRGAP2C has a distinct function in regulating synaptic development. The latter seems more plausible since SRGAP2B displays more CNVs in the human population than SRGAP2C (Dennis et al. 2012), strongly suggesting that these two paralogs have been under different positive selection pressures since their evolutionary emergence in the human population. Our data shows that SRGAP2C has a unique ability compared to SRGAP2B not only to delay synaptic maturation but also to increase synaptic density into adulthood. Considering that these proteins differ only in a small number of nonsynonymous base pair mutations, these results further underline the importance of the conversion of these five arginine residues during emergence of SRGAP2C.

Our results not only provide a molecular mechanism whereby SRGAP2B/C inhibits SRGAP2A function by targeting it for proteasome degradation but also, and for the first time, provide evidence that SRGAP2C (and its 5 specific arginine mutations) is more potent in its ability to inhibit synaptic maturation and maintain increased spine density throughout adulthood compared to SRGAP2B. This might explain why these two paralogs are under different selection pressures during recent human evolution and, more importantly, why emergence of SRGAP2C has played an important role during human brain evolution.

## Materials and methods

### Animals

All animals were handled according to protocols approved by the institutional animal care and use committee (IACUC) at Columbia University, New York. Timed pregnant CD1 mice (Crl:CD1(ICR)) were obtained from Charles River and maintained in a 12 h light/dark cycle. Juveniles correspond to mice at P21. Adults were P65 or older.

### Primary neuronal culture

Cortical neurons were *ex utero* electroporated (EUE) by injecting endotoxin-free DNA (500ng/µl for cell filler and 750ng/µl for each fusion protein construct) into the lateral ventricle of E15.5 mouse embryos and applying a current of 5 pulses of 20V for 100ms with 200ms intervals using a square wave electroporator (ECM 830, BTX). Cortices were subsequently dissected in complete HBSS (Hank’s Buffered Salt Solution (HBSS) supplemented with 2.5 mM Hepes (pH 7.4), 30 mM D-glucose, 1 mM CaCl_2_, 1mM MgSO_4_, and 4 mM NaHCO_3_). Dissociation was performed in papain (Worthington) supplemented with DNAse I (100 µg/µl) for 15 min at 37°C. We then performed three washes and triturated manually in plating medium. Cells were plated on cover slips or glass bottoms dishes coated with poly-D-lysine and cultured for 18-21 days *in vitro* (DIV). During the first 5 days, cells were cultured in neurobasal medium supplemented with 2.5% fetal bovine serum (FBS), B27, and L-glutamine. Half of the medium was replaced every 5 days with the same plating medium without FBS.

### Expression of proteins in cell lines and western blotting

Neuro-2a cell line was obtained from ATCC (CCL-131) and cultured according to the recommended protocol. Cells were transfected using Jet-Prime (Polyplus Transfection) according to the manufacturer protocol. After 48 h of transfection cells were scraped and lysed in lysis buffer (20mM HEPES, 150mM NaCl, 1mM EDTA, 1mM EGTA, 1% Triton-X100, cOmplete Protease Inhibitor Cocktail (Roche), Benzonase (EMD Millipore)) for 30 min at 4°C. Samples were prepared in Laemmli buffer (Bio-Rad) with 10% 2-Mercaptoethanol and boiled at 95°C for 5 min. Proteins were separated using SDS-PAGE and transferred to a polyvinylidene difluoride (PVDF) membrane (Immobilon-FL, EMD Millipore). Western blotting was performed using rabbit-anti-SRGAP2 N-terminal (raised against residues 193-205, 1:1000, [6]) and goat-anti-rabbit IgG conjugated to IRDye 600RD (1:20000, Li-Cor). Immunoblots were visualized on an Odyssey CLx Imaging System (Li-Cor).

### Fixed and live-cell imaging of dissociated neurons

For quantification of SRGA2A levels in soma and dendritic spines, neurons were fixed with 4% paraformaldehyde (Electron Microscopy Sciences) in PBS for 10 min at room temperature. Immunocytochemistry was performed for conditions where we used mTagBFP-HA-tagged SRGAP2A. Cells were incubated for 30 min in 0.2% Triton-X100 in PBS containing 5% goat serum to permeabilize and block nonspecific staining. Cells were incubated overnight at 4°C with mouse anti-HA primary antibody (1:1000, Anti-HA.11, Biolegend), washed 3× in PBS and incubated with Alexa-conjugated anti-mouse secondary antibody (1:500, Alexa Fluor 647, Invitrogen) for 1 h at room temperature. Coverslips were mounted on slides with Fluoromount-G aqueous mounting medium (ThermoFisher Scientific). Images were acquired on a Nikon A1 confocal microscope using a 60× (soma) or 100× (spines) objective. Exposure time and laser settings were optimized for bright cells to avoid pixel saturation and microscope settings were kept identical for each imaging session.

Live cell imaging was performed using a 40× objective on a Nikon A1 confocal microscope together with a stage incubation chamber (Tokai Hit) to maintain 37°C/5% CO_2_ culture conditions. Before imaging, medium was exchanged to recording medium consisting of pre-warmed cHBSS (Hank’s Buffered Salt Solution (HBSS) supplemented with 2.5 mM Hepes (pH 7.4), 30 mM D-glucose, 1 mM CaCl_2_, 1mM MgSO_4_, and 4 mM NaHCO_3_) with either 10 µM MG-132 or 200 µg/mL Cycloheximide (Sigma-Aldrich). Z-stacks that spanned the entire soma for each neuron were collected every 30 min for the indicated number of hours.

Analysis of images was performed using Nikon NIS-Elements (Nikon Corporation, Melville, NY). Background correction was performed by drawing an ROI at an area without fluorescent labeling and images were subsequently aligned to correct for sample drift. A z-stack maximum projection was generated for each neuron and using the filler signal ROIs were drawn over the soma, excluding the nucleus, or dendritic spines to measure fluorescent intensity.

### In utero cortical electroporation and in vivo analysis

*In utero* cortical electroporation was performed on timed pregnant CD1 females (Crl:CD1(ICR)). Endotoxin-free DNA containing 1µg/µl of each plasmid was injected into the ventricles of E15.5 embryos. Electroporation was performed using a square wave electroporator (ECM 830, BTX) by applying 5 pulses of 42 V for 50 ms with 500 ms intervals. When animals reached the indicated age they were anaesthetized with isoflurane and intracardiac perfusion with 4% paraformaldehyde (Electron Microscopy Sciences) in PBS was performed. Brains were subsequently isolated and incubated overnight in 4% paraformaldehyde/PBS at 4°C. Coronal brain sections were prepared by slicing brains at 100µm using a vibrating microtome (Leica VT1200S). Sections were mounted on glass slides in Fluoromount-G aqueous mounting medium (ThermoFisher Scientific).

Imaging of dendritic spines was performed on optically isolated neurons using a Nikon A1 confocal microscope and 100× objective. Using Nikon NIS-Elements software maximum intensity projections were generated of oblique dendrites originating from the main apical trunk. Spine density and head size was quantified by using the Venus filler signal to draw ROIs around the spine head.

### Statistical analysis

A minimum of three independent experiments were performed unless otherwise stated. For *in vivo* spine analysis, data was obtained from at least six animals from a minimum of two independent litters. Statistical analysis was performed using Prism (Graphpad Software). Normality was checked using D’Agostino-Pearson omnibus normality test. A non-parametric test (Mann-Whitney) was used when distribution deviated significantly from normality. A test was considered significant when p < 0.05.

## Acknowledgements

We thank Virginie Courchet, Miyako Hirabayashi and Qiaolian Liu for excellent technical help. We thank members of the Polleux lab for valuable discussions and inputs. This work was supported by NIH (RO1NS067557) (FP), the Roger De Spoelberch Foundation (FP), Netherlands Organization for Scientific Research (NWO Rubicon 825.14.017) (ERES), the European Molecular Biology Organization (EMBO Long-Term Fellowship ALTF 1055-2014) (ERES) and K99.

---

**S1 Fig.**
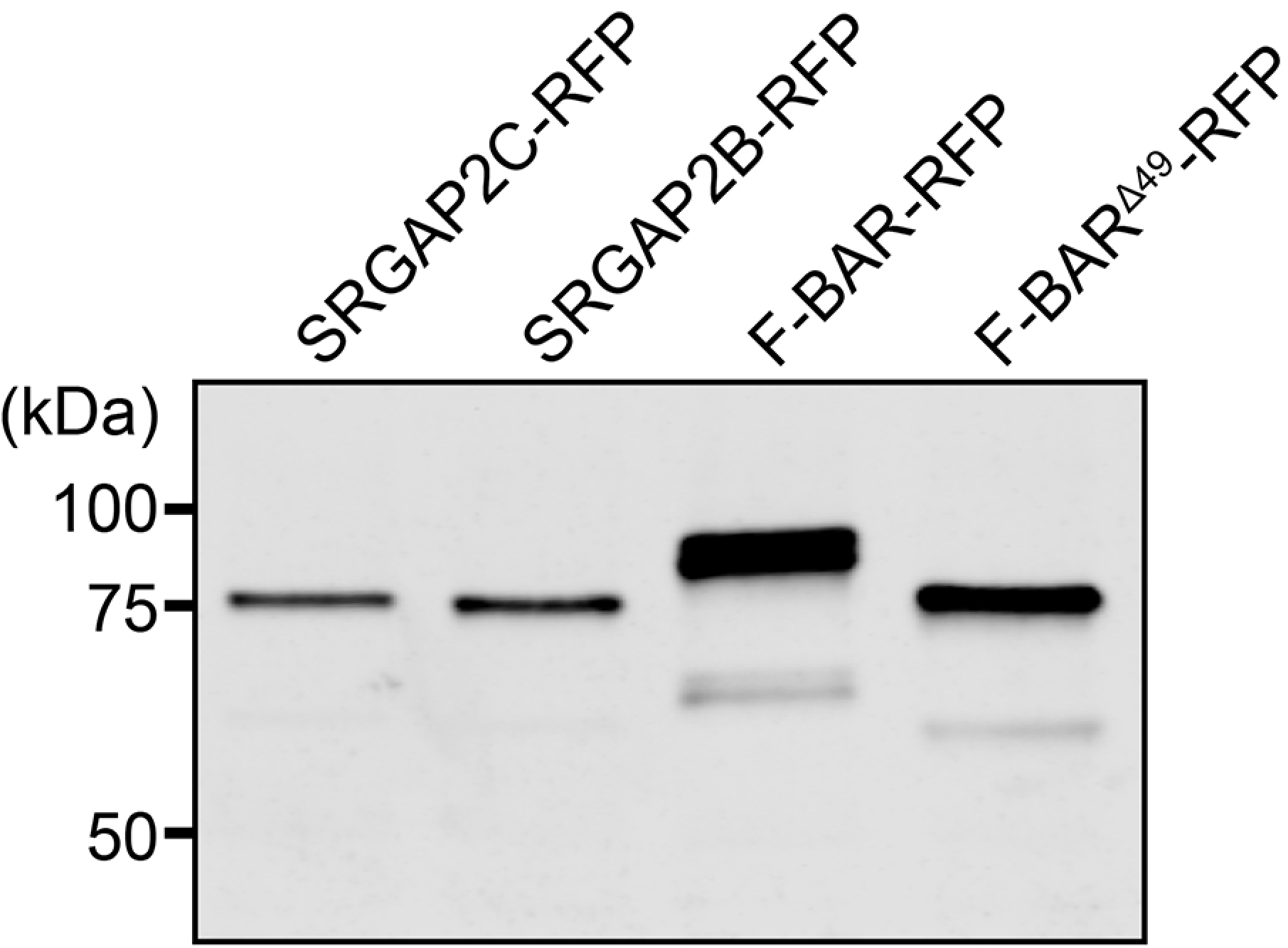
Testing expression of fusion proteins in Neuro-2a cells. Western blot showing expression of indicated mRFP-fusion proteins in Neuro-2a cell lines. SRGAP2C-RFP, SRGAP2B-RFP, and F-BAR^Δ49^-RFP run slightly lower than F-BAR-RFP due to their lack of the last C-terminal 49 amino acids.

**S2 Fig.**
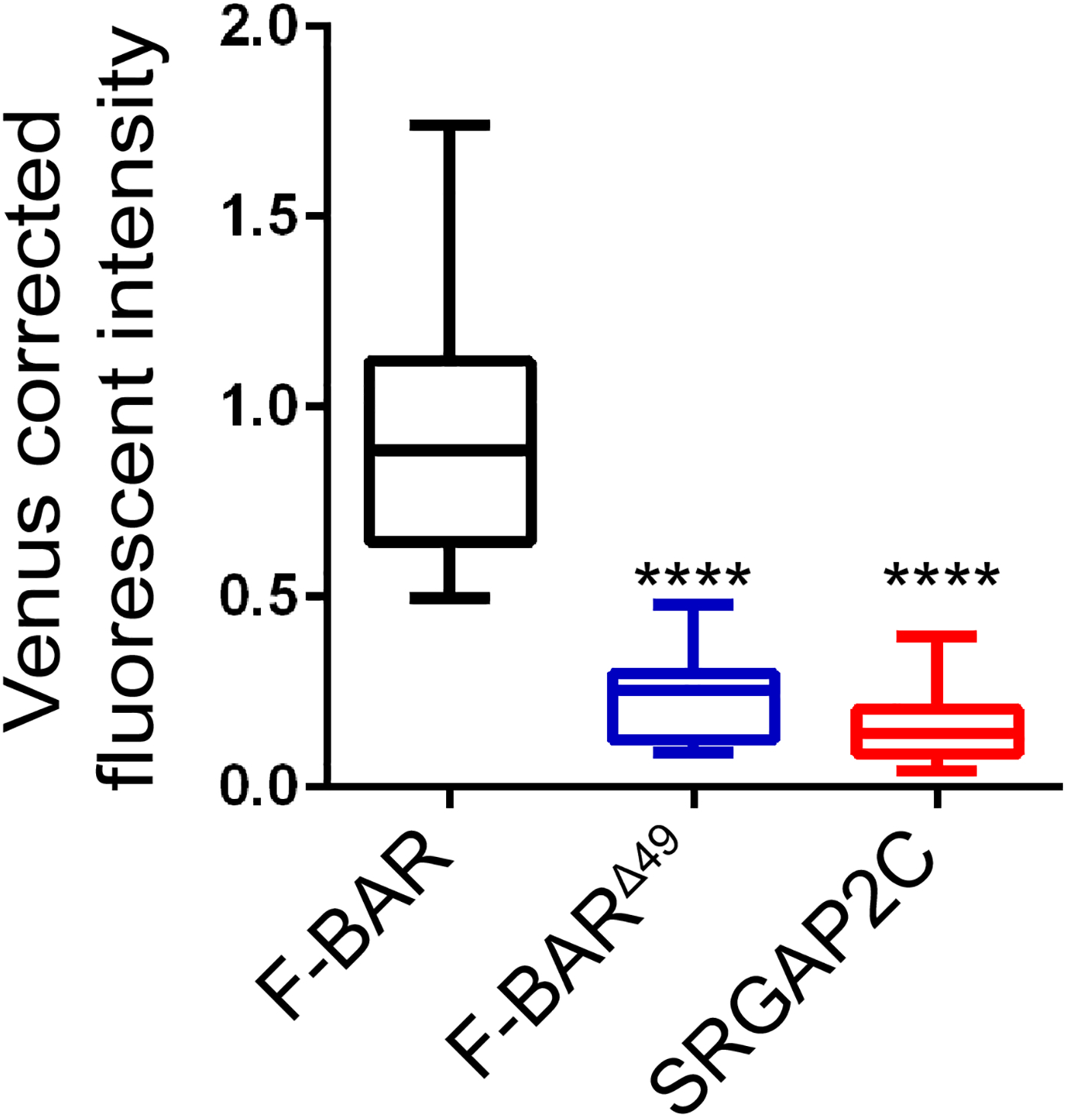
Expression levels for F-BAR proteins in cortical pyramidal neurons. Box plots showing quantification of mRFP-tagged F-BAR, F-BAR^Δ49^, and SRGAP2C fluorescent intensity in the soma of cultured cortical pyramidal neurons. n_F-BAR_ = 21, n_F-BAR_^Δ49^ = 33, n_SRGAP2C_ = 31; ****p < 0.0001.

**S3 Fig.**
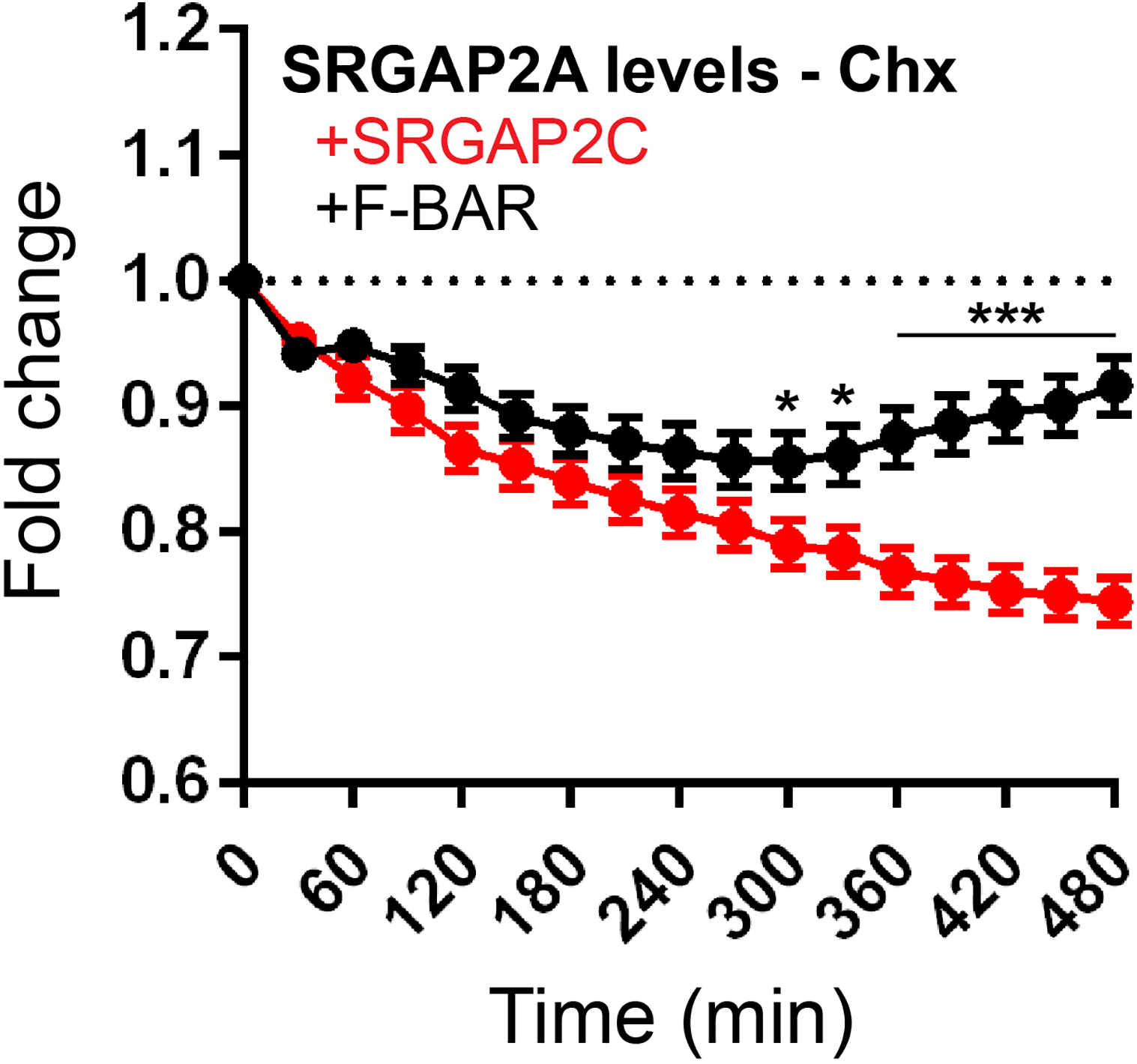
SRGAP2A turnover when co-expressed with SRGAP2C. Quantification of SRGAP2A fluorescent intensity upon treatment with cycloheximide (Chx). When co-expressed with SRGAP2C, the reduction of SRGAP2A levels is significantly increased compared to co-expression with F-BAR. n_F-BAR_ = 24, n_SRGAP2C_ = 29; *p < 0.05, ***p < 0.001; mean ± SEM.

**S4 Fig.**
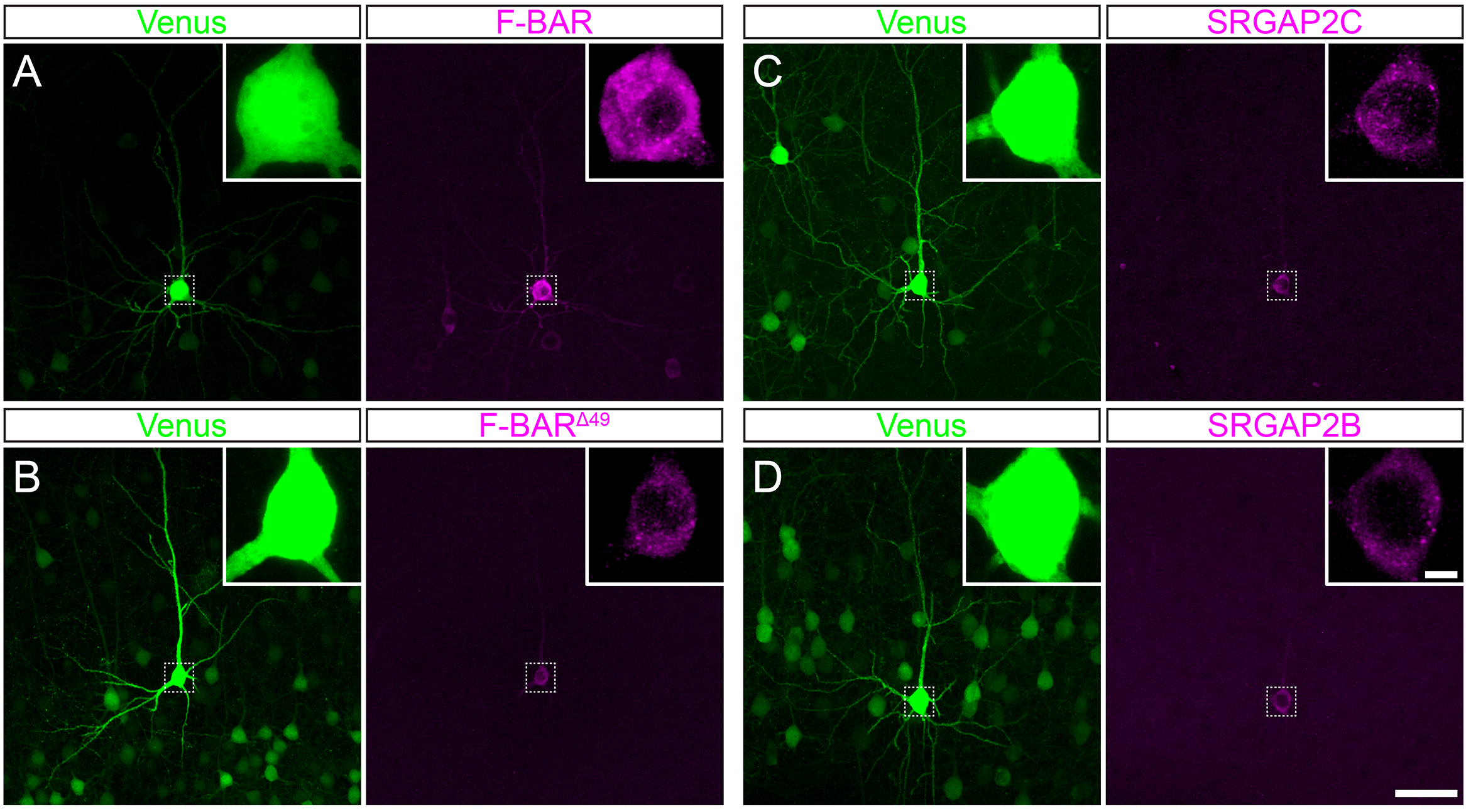
*In vivo* expression of F-BAR proteins in layer 2/3 cortical pyramidal neurons. Representative images showing *in utero* electroporated layer 2/3 cortical pyramidal neurons expressing Venus with either mRFP-tagged F-BAR, F-BAR^Δ49^, SRGAP2B, or SRGAP2C. Insets show high magnification images of soma (indicated with dashed rectangle). Scale bars: 50 µm, and 5 µm for inset.

## References

1. Sousa AMM, Meyer KA, Santpere G, Gulden FO, Sestan N. Evolution of the Human Nervous System Function, Structure, and Development. Cell. 2017;170: 226–247. doi:10.1016/j.cell.2017.06.036

2. Suzuki IK, Gacquer D, Van Heurck R, Kumar D, Wojno M, Bilheu A, et al. Human-Specific NOTCH2NL Genes Expand Cortical Neurogenesis through Delta/Notch Regulation. Cell. 2018; 1370–1384. doi:10.1016/j.cell.2018.03.067

3. Fiddes IT, Lodewijk GA, Mooring M, Bosworth CM, Ewing AD, Mantalas GL, et al. Human-Specific NOTCH2NL Genes Affect Notch Signaling and Cortical Neurogenesis. Cell. Elsevier; 2018;173: 1356–1369.e22. doi:10.1016/j.cell.2018.03.051

4. Florio M, Albert M, Taverna E, Namba T, Brandl H, Lewitus E, et al. Human-specific gene ARHGAP11B promotes basal progenitor amplification and neocortex expansion. Science. 2015;347: 1465–1470. doi:10.1126/science.aaa1975

5. Florio M, Heide M, Pinson A, Brandl H, Albert M, Winkler S, et al. Evolution and cell-type specificity of human-specific genes preferentially expressed in progenitors of fetal neocortex. Elife. 2018;7: 1–37. doi:10.7554/eLife.32332

6. Charrier C, Joshi K, Coutinho-Budd J, Kim J-E, Lambert N, de Marchena J, et al. Inhibition of SRGAP2 function by its human-specific paralogs induces neoteny during spine maturation. Cell. 2012;149: 923–35. doi:10.1016/j.cell.2012.03.034

7. Fossati M, Pizzarelli R, Schmidt ER, Kupferman J V, Stroebel D, Polleux F, et al. SRGAP2 and Its Human-Specific Paralog Co-Regulate the Development of Excitatory and Inhibitory Synapses. Neuron. 2016;91: 356–369. doi:10.1016/j.neuron.2016.06.013

8. Dennis MY, Nuttle X, Sudmant PH, Antonacci F, Graves TA, Nefedov M, et al. Evolution of human-specific neural SRGAP2 genes by incomplete segmental duplication. Cell. Elsevier Inc.; 2012;149: 912–22. doi:10.1016/j.cell.2012.03.033

9. Ataman B, Boulting GL, Harmin DA, Yang MG, Baker-Salisbury M, Yap EL, et al. Evolution of Osteocrin as an activity-regulated factor in the primate brain. Nature. Nature Publishing Group; 2016;539: 242–247. doi:10.1038/nature20111

10. Enard W, Gehre S, Hammerschmidt K, Hölter SM, Blass T, Somel M, et al. A Humanized Version of Foxp2 Affects Cortico-Basal Ganglia Circuits in Mice. Cell. 2009;137: 961–971. doi:10.1016/j.cell.2009.03.041

11. Boyd JL, Skove SL, Rouanet JP, Pilaz LJ, Bepler T, Gordân R, et al. Human-chimpanzee differences in a FZD8 enhancer alter cell-cycle dynamics in the developing neocortex. Curr Biol. 2015;25: 772–779. doi:10.1016/j.cub.2015.01.041

12. Guerrier S, Coutinho-Budd J, Sassa T, Gresset A, Jordan NV, Chen K, et al. The F-BAR domain of srGAP2 induces membrane protrusions required for neuronal migration and morphogenesis. Cell. 2009;138: 990–1004. doi:10.1016/j.cell.2009.06.047

13. Bacon C, Endris V, Rappold G. Dynamic expression of the Slit-Robo GTPase activating protein genes during development of the murine nervous system. J Comp Neurol. 2009;513: 224–36. doi:10.1002/cne.21955

14. Sudmant PH, Kitzman JO, Antonacci F, Alkan C, Malig M, Tsalenko A, et al. Diversity of human copy number variation and multicopy genes. Science. 2010;330: 641–6. doi:10.1126/science.1197005

15. Sporny M, Guez-Haddad J, Kreusch A, Shakartzi S, Neznansky A, Cross A, et al. Structural History of Human SRGAP2 Proteins. Mol Biol Evol. 2017;34: 1463–1478. doi:10.1093/molbev/msx094

16. Guez-Haddad J, Sporny M, Sasson Y, Gevorkyan-Airapetov L, Lahav-Mankovski N, Margulies D, et al. The Neuronal Migration Factor srGAP2 Achieves Specificity in Ligand Binding through a Two-Component Molecular Mechanism. Structure. Elsevier Ltd; 2015; 1–12. doi:10.1016/j.str.2015.08.009

17. Okada H, Uezu A, Mason FM, Soderblom EJ, Moseley MA, Soderling SH. SH3 domain-based phototrapping in living cells reveals Rho family GAP signaling complexes. Sci Signal. 2011;4: rs13. doi:10.1126/scisignal.2002189

18. Harris KM, Stevens JK. Dendritic spines of CA 1 pyramidal cells in the rat hippocampus: serial electron microscopy with reference to their biophysical characteristics. J Neurosci. 1989;9: 2982–2997. doi:10.1021/la0512603

19. Arellano JI. Ultrastructure of dendritic spines: correlation between synaptic and spine morphologies. Front Neurosci. 2007;1: 131–143. doi:10.3389/neuro.01.1.1.010.2007

20. Matsuzaki M, Ellis-Davies GCR, Nemoto T, Miyashita Y, Iino M, Kasai H. Dendritic spine geometry is critical for AMPA receptor expression in hippocampal CA1 pyramidal neurons. Nat Neurosci. 2001;4: 1086–1092. doi:10.1038/nn736

21. Dennis MY, Harshman L, Nelson BJ, Penn O, Cantsilieris S, Huddleston J, et al. The evolution and population diversity of human-specific segmental duplications. Nat Ecol Evol. 2017;1: 0069. doi:10.1038/s41559-016-0069

22. Dennis MY, Eichler EE. Human adaptation and evolution by segmental duplication. Curr Opin Genet Dev. Elsevier Ltd; 2016;41: 44–52. doi:10.1016/j.gde.2016.08.001

23. Dougherty ML, Nuttle X, Penn O, Nelson BJ, Huddleston J, Baker C, et al. The birth of a human-specific neural gene by incomplete duplication and gene fusion. Genome Biol. Genome Biology; 2017;18: 1–16. doi:10.1186/s13059-017-1163-9

24. Franchini LF, Pollard KS. Human evolution: The non-coding revolution. BMC Biol. BMC Biology; 2017;15: 1–12. doi:10.1186/s12915-017-0428-9

